# RHD6LA regulates root hair responses to both symbionts and commensals

**DOI:** 10.1101/2025.05.22.655471

**Authors:** Francesca Tedeschi, Johan Quilbé, Lavinia Ioana Fechete, Sofie Jin Vistisen Christiansen, Stig Uggerhøj Andersen

## Abstract

While intracellular symbiosis with rhizobia relies on Nod factor signaling through the conserved common symbiosis signaling pathway (CSSP), it remains unclear how legumes simultaneously manage interactions with commensal soil microbes. Using single cell RNA-sequencing, we show that commensal soil bacteria induce a Nod factor-independent transcriptional response in specific root hairs. This response is similar to the rhizobium response in the CSSP-deficient *cyclops* mutant, which is unable to accommodate rhizobia in root hair infection threads. Both responses include the nodulation gene *NSP2* and a transcription factor, which we name *ROOT HAIR DEFECTIVE 6 LIKE A* (*RHD6LA*). We show that *RHD6LA* is required for facilitating infection thread formation in response to rhizobia and for preventing exaggerated root hair responses to commensal soil bacteria. The overlap between commensal and symbiotic signaling highlights the complexity of legume-microbe interactions at the root hair interface and suggests new mechanisms for microbial discrimination in rhizobium-responsive root hairs.

## Main

Plant roots harbor a diverse community of bacteria known as the root microbiota, which play an important role in shaping plant growth, health, and overall ecosystem function ^1, 2, 3^ ^4^. This association between plants and their root-associated bacteria has gained increasing attention due to its potential implications for agriculture and environmental sustainability ^5, 6^ ^7^. However, despite the importance of the root microbiota, the understanding of the mechanisms that govern plant-microbe interactions to sculpt and establish bacterial communities around roots remains rudimentary at the molecular level.

The exception is the symbiotic interaction between legumes and rhizobia where specific molecular pathways and genetic components critical for successful establishment are well defined. The common symbiosis signaling pathway (CSSP) in legumes is essential for establishing long-term, intracellular associations with both nitrogen-fixing rhizobia and arbuscular mycorrhizal (AM) fungi ^8^. Recognition of Nod- or Myc-factors triggers specific signaling responses, leading to characteristic calcium spiking patterns that are interpreted by CCAMK ^9^. These calcium signals subsequently activate *CYCLOPS*, a transcriptional regulator positioned at the interface between mycorrhizal and rhizobial symbioses ^10^. *CYCLOPS*, in turn initiates the transcription of several downstream symbiotic genes, including *NSP1* and *NSP2*, which are essential for nodulation and also promote colonization by AM fungi ^11^. In *cyclops* mutants, the early Nod factor perception and Nod factor receptor-mediated signaling events in root hairs, including calcium spiking, remain functional. This allows initial root hair curling and rhizobial entrapment. However, downstream activation of the infection thread (IT) program is impaired, resulting in rhizobia accumulating in infection pockets without successful IT progression ^10^. Despite this block in epidermal infection, cortical responses are still activated, leading to initiation of nodule organogenesis in the absence of infection threads ^10^.

Recent single cell RNA-sequencing analyses have contributed to elucidation of the cellular expression patterns of symbiotic genes, revealing specific transcriptional profiles within root hair cells that directly interface with rhizobia ^12^. Application of similar approaches to the study of root-microbiome interactions is difficult because of the absence of established marker genes for the colonization of the root surface or the entry of commensal bacteria into host tissues. While several studies have examined plant expression responses to mono-inoculation with bacterial strains possessing specific characteristics ^13^ ^14^ ^15^ ^16^ ^17^ or within the context of complex microbial communities, these investigations predominantly highlighted broad biological processes, including stress responses, immunity, cell wall modification, nutrient uptake, primary metabolism, and epigenetic regulation ^18^ ^19^ ^20^ ^21^ .

Here, we use single cell RNA-sequencing of *Lotus japonicus* roots inoculated with a bacterial Synthetic Community (SynCom) with or without rhizobia to compare plant responses to symbiotic and commensal bacteria at the cellular level.

## Results

### A bacterial SynCom enriched in plant interaction-related genes

To establish our commensal inoculant, we selected 19 fully sequenced strains (SynCom19) from the *Lj*-SPHERE culture collection ^17^. We based the strain selection on their content of genes potentially associated with plant interactions and on their rhizosphere colonization characteristics in a diverse panel of *Lotus japonicus* (Lotus) accessions ^22^. SynCom19 includes strains from eight distinct bacterial families **(Supplemental Fig. 1**). Many of the strains carry genes associated with plant growth-promoting (PGP) functions, including cytokinin production (miaABE) ^23^, ethylene precursor degradation (acdS) ^24^, phosphate mobilization and solubilization (phyC, gcd, pqqBCDEG) ^25^, spermidine production (speBE) ^26^, quercetin degradation (Qdx) ^27^, antagonism (phlG)^28^, and mineral uptake (cysA) ^29^. We characterized the taxonomy of the strains using phylogenetic trees based on full-length *recA* sequences and, where possible, classified strains to the species level through whole-genome alignment with reference genomes (**Supplemental Table 1**). SynCom19 does not include compatible symbionts capable of nodulating Lotus, allowing us to specifically study plant interactions with a community of commensal bacteria.

### Pervasive root transcriptional response to commensal bacteria

To categorize cells based on tissue type and investigate their responses to commensal bacteria alone and to commensal bacteria in combination with the Lotus symbiont *M. loti* R7A (R7A), we carried out scRNA-seq of protoplasts from Lotus wild-type roots 5 dpi with SynCom19, SynCom19+R7A and mock-inoculated. We characterized a total of 50,996 cells (**Source Data “SynCom5dpi_Cells”)** with a median of 2,101 unique molecular identifiers and ∼1,200 transcripts per cell after filtering. After integration, the samples were clustered using Seurat^30^ (**Fig. 1a**). Cellular identities of individual clusters were assigned using homologous markers from Arabidopsis, marker gene information from Lotus Base^31^ and promoter-reporter lines as previously reported^32^.

**Figure 1.**
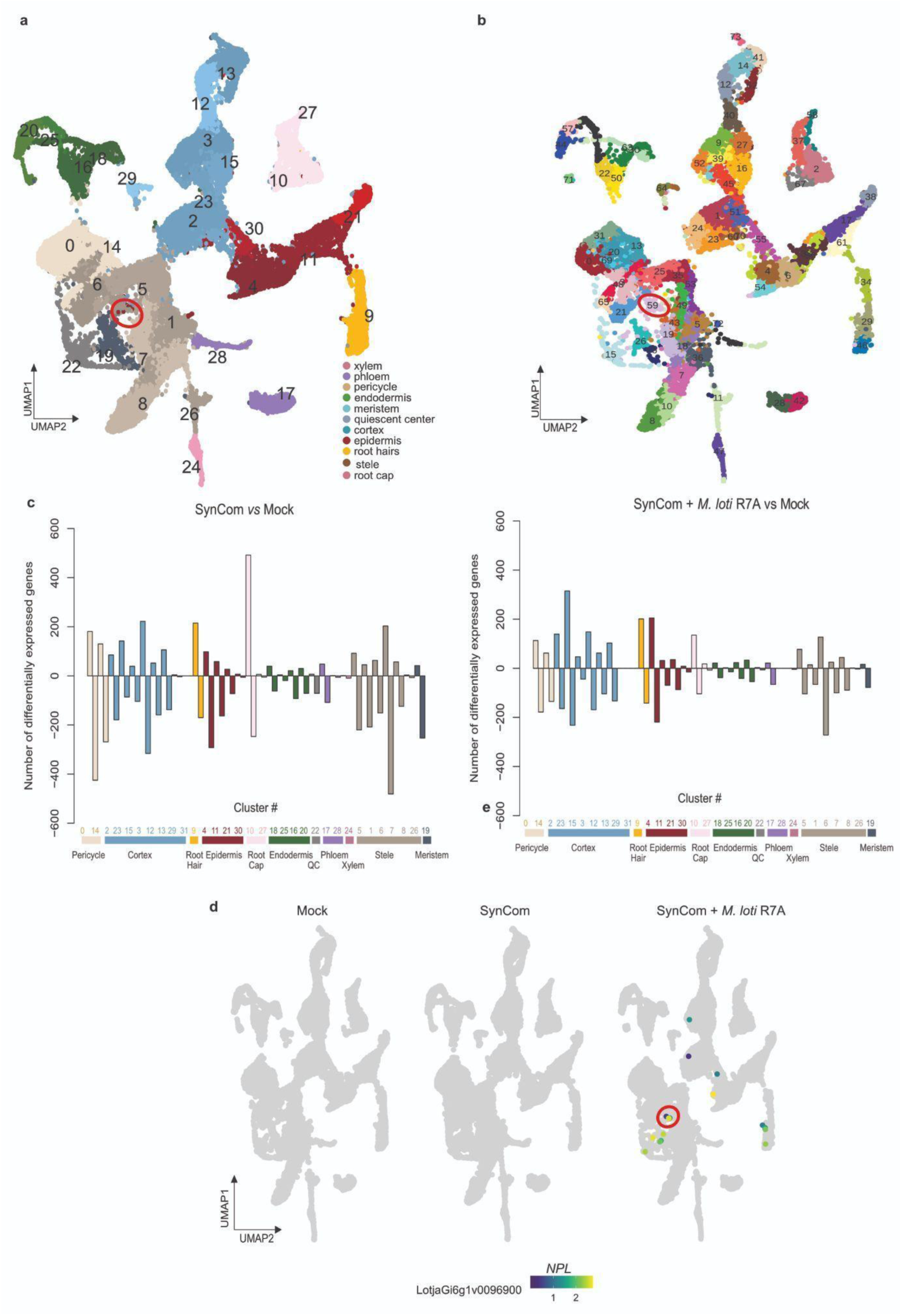
A cellular atlas of Lotus roots inoculated with SynCom +/- *M. loti* R7A. **a**) UMAP of control, SynCom and SynCom + *M. loti* R7A inoculated root cells 5dpi showing clusters for known root cell types. **b**) The UMAP reclustering using the cluster numbers created with TopOMetry ^34^ allowed the assignment of the cluster containing marker genes of rhizobia responding cells. **c**) DEGs in the clusters in response to SynCom presence compared to mock (left) and DEGs in the clusters in response to SynCom + *M. loti* R7A compared to mock (right). Complete lists of differentially expressed genes and marker genes can be found in Source Data. **d**) Example of symbiotic specific gene induced in presence of *M. loti* (NPL).

Using MAST^33^ to conduct differential gene expression analysis, we found distinct responses of different cell types to inoculation with SynCom19 at 5 dpi compared to the control (**Source Data “SynCom5dpi_DE_Genes” and Fig. 1c**). Most cell types exhibited a higher number of downregulated than upregulated genes, but root cap and root hair cells showed the reverse trend. The least responsive cell types were xylem, endodermis and phloem. Root cap, root hair, pericycle, stele, and meristem had the highest total number of DEGs (**Fig. 1c).** Expression patterns of all genes can be browsed online using our ShinyApp: https://lotussinglecell.shinyapps.io/lotus_japonicus_single_cell_microbiome/.

To compare the plant root response to SynCom19 in the presence and absence of rhizobia, we first examined DEGs for the SynCom19+R7A versus Mock samples (**Source Data “SynComR7A_5dpi_DE_Genes”**). We identified rhizobial infection and nodulation-related genes, including *NPL*, which were specifically induced in root hairs and cortical cells following inoculation with SynCom19+R7A (**Fig. 1d**). The infected cortical cells did not form a distinct cluster using Seurat’s default dimensionality reduction function (**Fig. 2a,d)**. In contrast, clustering using TopOmetry ^34^ produced a single and well-defined cluster containing the infected cortical cells (**Fig. 1b**). This cluster was also represented in the Mock and SynCom19 samples, but did not show *NPL* expression (**Fig. 1d**), indicating that there was no R7A contamination and that Nod-factor signaling was exclusively active in the SynCom19+R7A samples.

**Figure 2.**
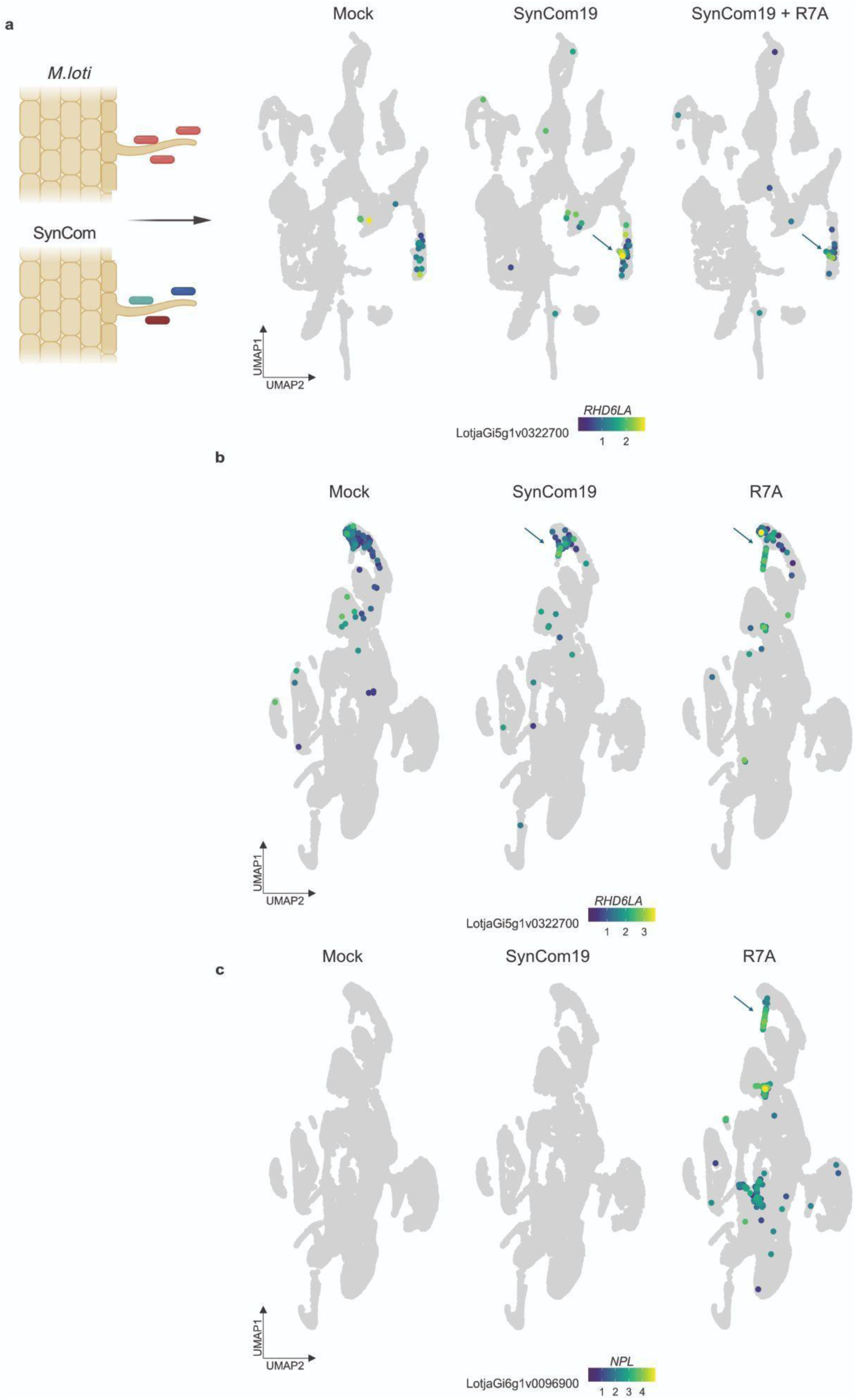
*ROOT HAIR DEFECTIVE 6 LIKE A* is induced in root hairs upon bacteria inoculation. **a**) A defined region in the root hair (indicated by the blue arrow) exhibits *RHD6LA* gene induction in response to both SynCom19 and SynCom19 + R7A treatments. **b**-**c**) The current SynCom19 data integrated with previously published datasets ^32^, one featuring Gifu WT samples mock-inoculated and R7A-inoculated at 10 dpi and another including Gifu WT and *cyclops* samples that were mock-inoculated and R7A-inoculated at 5 dpi. **b**) Induction of *RHD6LA* in a subset of root hairs observed only in samples inoculated with SynCom19 and R7A. **c**) Expression of the rhizobium infection marker *NPL*. Note that in both (b) and (c), Mock and R7A and samples represent combined Gifu 5 and 10 dpi data, but do not include *cyclops* data.

### Overlapping root hair transcriptional responses to symbionts and commensals

Although classical rhizobium infection markers like *NPL* were not induced by SynCom19 alone, we examined the extent to which transcriptional responses to symbiotic and commensal bacteria overlapped. To identify genes induced by both R7A alone and SynCom19 without R7A, we compared the SynCom19 DEGs to R7A DEGs from a previous study ^32^. In addition, we integrated our current data with previously published 5 and 10 dpi *M. loti* R7A scRNA-seq data ^32^ to allow direct comparison of cell populations across studies (**Fig. 2**).

We identified a set of 23 genes (**Source Data “Common_genes”**) that were induced by both SynCom19 and R7A. Among these genes, *LotjaGi5g1v0322700* exhibited a distinct induction pattern in a subset of rhizobium-responsive root hairs (**Fig. 2**). *LotjaGi5g1v0322700* encodes a bHLH transcription factor, which is a close homolog of the Arabidopsis *AT1G27740* gene known as *ROOT HAIR DEFECTIVE 6-LIKE 4 (RSL4)* and we named it *ROOT HAIR DEFECTIVE 6 LIKE A* (*RHD6LA*).

The SynCom19 commensals do not activate Nod factor signaling, whereas R7A does (**Fig. 1d**). In contrast, both treatments induce *RHD6LA* in root hairs (**Fig. 2b**). To further investigate the root hair response to commensals exemplified by *RHD6LA*, we re-clustered Mock and SynCom19 root hair cells, resulting in ten subclusters (**Fig. 3a**). Among these, only subcluster 7, comprising 92 out of 1658 root hair cells, consistently expressed *RHD6LA*. This subcluster represents the SynCom19-responsive root hair cells, and we will refer to it as SynCom19_RHsubcluster7 (**Fig. 3b**). By comparison with the remaining root hair cells, we identified 227 marker genes for SynCom19_RHsubcluster7 (**Fig. 3a-b**, **Source Data “SynCom19_RHsubcluster7”**).

**Figure 3.**
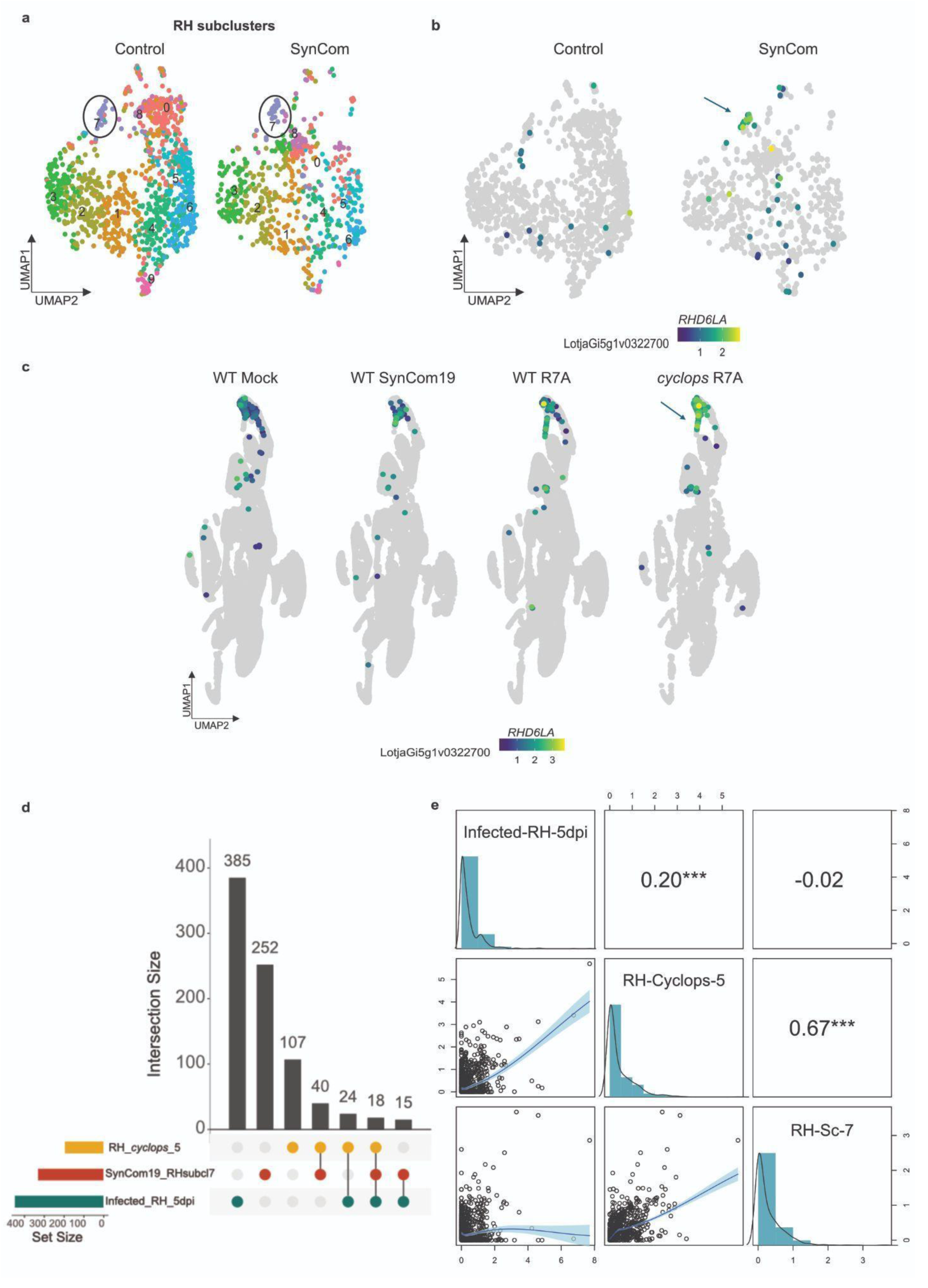
Similarities between wild-type SynCom19 and *cyclops* R7A responses. **a**) UMAP of reclustered root hair cells from control and SynCom19 samples. **b**) Subcluster 7 showed strong and specific induction of *RHD6LA* on SynCom19 inoculation. **c**) A specific root hair population shows a high induction of *RHD6LA* in *cyclops* upon *M. loti* R7A inoculation. **d**)Upset plot showing the markers for *cyclops* R7A responding root hairs (RH *cyclops*), SynCom19 root hair subcluster 7 (SynCom19_RHsubcl7), Infected RH 5 dpi markers ^32^ and their overlaps (pct.2 <= 0.1). e) Bivariate scatter plots, histograms, and Spearman correlation values for the log2 average expression of 841 markers in three cell populations *cyclops* R7A responding root hairs (RH-*Cyclops-5*), SynCom19 root hair subcluster 7 (RH-Sc-7), and Infected RH 5 dpi.

### Wild-type SynCom19 and *cyclops* R7A root hair responses are similar

Focusing on SynCom19 and R7A responsive root hairs across our integrated object, which comprises data from Mock, SynCom19, SynCom19+R7A and R7A treated wild-type plants and Mock and R7A treated *cyclops* mutants, we observed pronounced *RHD6LA* induction by R7A in *cyclops* root hairs (**Fig. 3c**). Since Nod factor signalling is absent in SynCom19 and only partly active in *cyclops* mutants ^10^, we asked if the wild-type SynCom19 root hair response was more similar to the wild-type or to the *cyclops* R7A root hair response. To address this question, we focused on three root hair subpopulations: Infected_RH_5dpi — root hairs responding to R7A at 5 dpi, SynCom19_RHsubcluster7 — the subset of SynCom19-responsive root hairs at 5 dpi identified in this study, and RH_*cyclops*_5 — a root hair population from *cyclops* mutant roots at 5 dpi showing an aberrant response to R7A. We then compared these populations to the corresponding mock-inoculated root hair cells at 5dpi, identifying 325 marker genes for SynCom19_RHsubcluster7, 422 for Infected_RH_5dpi, and 149 for RH_*cyclops*_5 cells, totaling 841 unique genes (**Fig. 3d**). To assess the transcriptional similarity across the three root hair cell populations, we compared the pseudobulk expression levels of the 841 differentially expressed genes for each population, revealing a significantly higher correlation between SynCom19_RHsubcluster7 and RH_*cyclops*_5 root hair expression profiles than between SynCom19_RHsubcluster7 and Infected_RH_5dpi cell (*P* < 0.0001, Pearson and Filon’s z (1898) test) (**Fig. 3e**), a pattern also observed at the level of individual gene expression (**Supplementary fig. 2**). In addition, the correlation between RH_*cyclops*_5 and SynCom19_RHsubcluster7 cells was higher than that between RH_*cyclops*_5 and Infected_RH_5dpi (*P* < 0.0001, Pearson and Filon’s z (1898) test), although both these populations were inoculated with R7A (**Fig. 3e**). SynCom19 thus induces a transcriptional state in root hair cells that closely resembles that of R7A-treated *cyclops* mutants, which is characterized by activation of genes associated with early root hair responses but lacks infection thread progression ^10^.

### *RHD6L*A controls root hair response to commensals and symbionts

We identified 18 overlapping genes between the SynCom19_RHsubcluster7, Infected_RH_5dpi and RH_*cyclops*_5 markers (**Fig. 3d, Source data “Common_subcl7_cyclops_5INF”**). In addition to induction during a compatible symbiotic interaction, these were all induced in the absence of Nod factor by SynCom19 and by R7A in the absence of the CSSP transcription factor CYCLOPS, suggesting that they may be components of a general bacterial perception system in root hairs. *RHD6LA* was included in this set of 18 genes, but its function has not been described. Since *RHD6LA* was induced by both symbionts and commensals, it could potentially play a role in both interactions. Legume roots respond to the presence of symbiotic rhizobia through the curling of root hairs to facilitate bacterial attachment and subsequent infection in a Nod factor-dependent manner ^40^. Given that the same population of root hairs exhibited a transcriptional response to SynCom19 (**Fig. 2b** and **Supplemental Fig. 1b**), we investigated if SynCom19 also elicits a morphological response in root hairs. To assess this and explore the potential involvement of *RHD6LA*, we obtained two LORE1 *rhd6la* insertion lines ^41^ and examined root hair morphology in wild-type and *rhd6la* mutant plants at 5 dpi with SynCom19 using confocal microscopy. In both cases, most root hairs did not respond to the treatment, but displayed a normal elongated and tubular shape (**Fig. 4a**). Some wild-type and *rhd6la* mutant root hairs, however, exhibited clear deformation upon inoculation with SynCom19 (**Figure 4b-c**). The swollen and other abnormal root hair phenotypes observed in the presence of commensals were significantly more frequent in the *rhd6la* mutants, indicating that *RHD6LA* plays a role in regulating root hair responses to commensal bacteria (**Fig. 4d**). To determine if *RHD6LA* also plays a role in symbiotic interactions, we compared wild-type and *rhd6la* infection thread (10 dpi) and nodule (21 dpi) counts. Consistent with a compromised symbiotic response, comparison with WT (n=12) revealed a significant reduction in *rhd6la-1* and *rhd6la-2* for infection thread density (n=12) and nodule count (**Figure 4e-f**). Consistent with its transcriptional profile, *RHD6LA* thus regulates root hair responses to both commensal and symbiotic bacteria.

**Figure 4.**
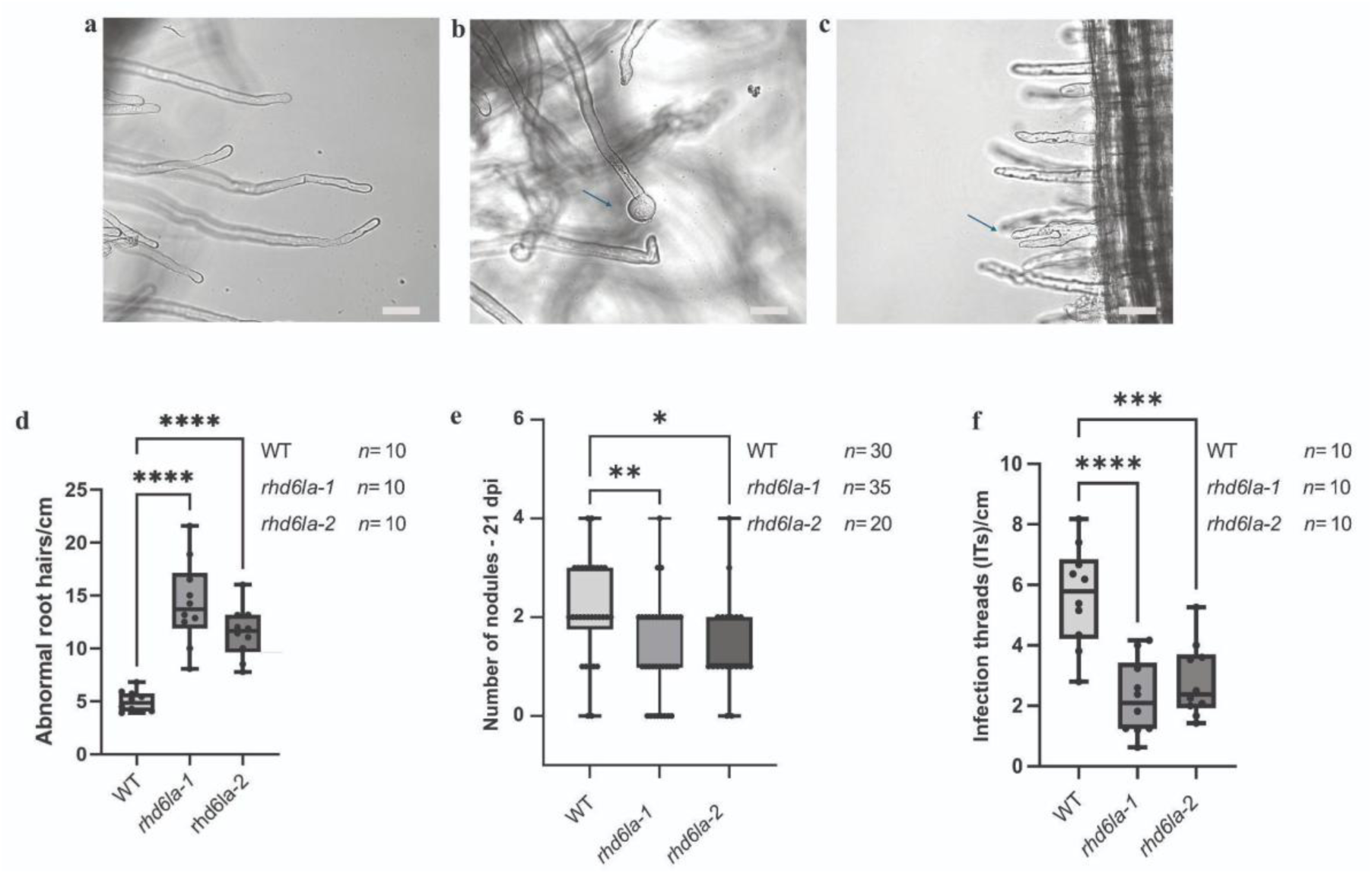
Responses to SynCom19 in *rhd6la* mutants. Panel a)-c) Confocal pictures of non-responsive (**a**) and abnormal *rhd6la* root hair cells (**b-c**) in the presence of SynCom19 5dpi. **d**) Ten entire roots were examined, and the phenotypes of wild-type and mutant root hairs were scored for abnormalities. **e**) Nodulation assay of wild-type (n= 30) and mutant plants (*rhd6la-*1 n= 35; *rhd6la-2* n= 20) inoculated with *M. loti* R7A 21 dpi. **f**) IT counting of wild-type (n= 10) and *rhd6la* mutant alleles (n=10) plants 10 dpi *P < 0.05, **P < 0.01, ***P < 0.001, ****P < 0.0001 (Dunnett’s test). Bars, 50 µM.

## Discussion

scRNA-seq studies have provided significant insights into the transcriptional regulation of symbiotic signaling in legume-rhizobia interactions, highlighting the complexity and specificity of gene expression in different cell types and stages of symbiosis^12^. While commensals are known to interact with root systems and potentially influence plant health and growth ^17,18^, the transcriptional responses and signaling pathways involved in these interactions have not been thoroughly characterized.

Our study demonstrates that plant roots mount a substantial transcriptional response to commensal bacteria. It differs between cell types and includes a specific response in a population of rhizobium-responsive root hairs. CYCLOPS is well known for its essential role in intracellular infection of plant roots by rhizobia and arbuscular mycorrhizal fungi ^10^ ^43^. In *cyclops* mutants, Nod factor perception remains intact, allowing root hair curling and initiation of nodule primordia, but rhizobial infection is abrogated at an early stage, leaving rhizobia trapped in curled root hairs ^10^ (**Fig. 5**). We observed that commensal bacteria, which do not produce Nod factors, induce a transcriptional response in wild-type root hairs that overlaps significantly with the response to Nod factor-producing rhizobia in *cyclops* mutant root hairs (**Fig. 3e**).

**Figure 5.**
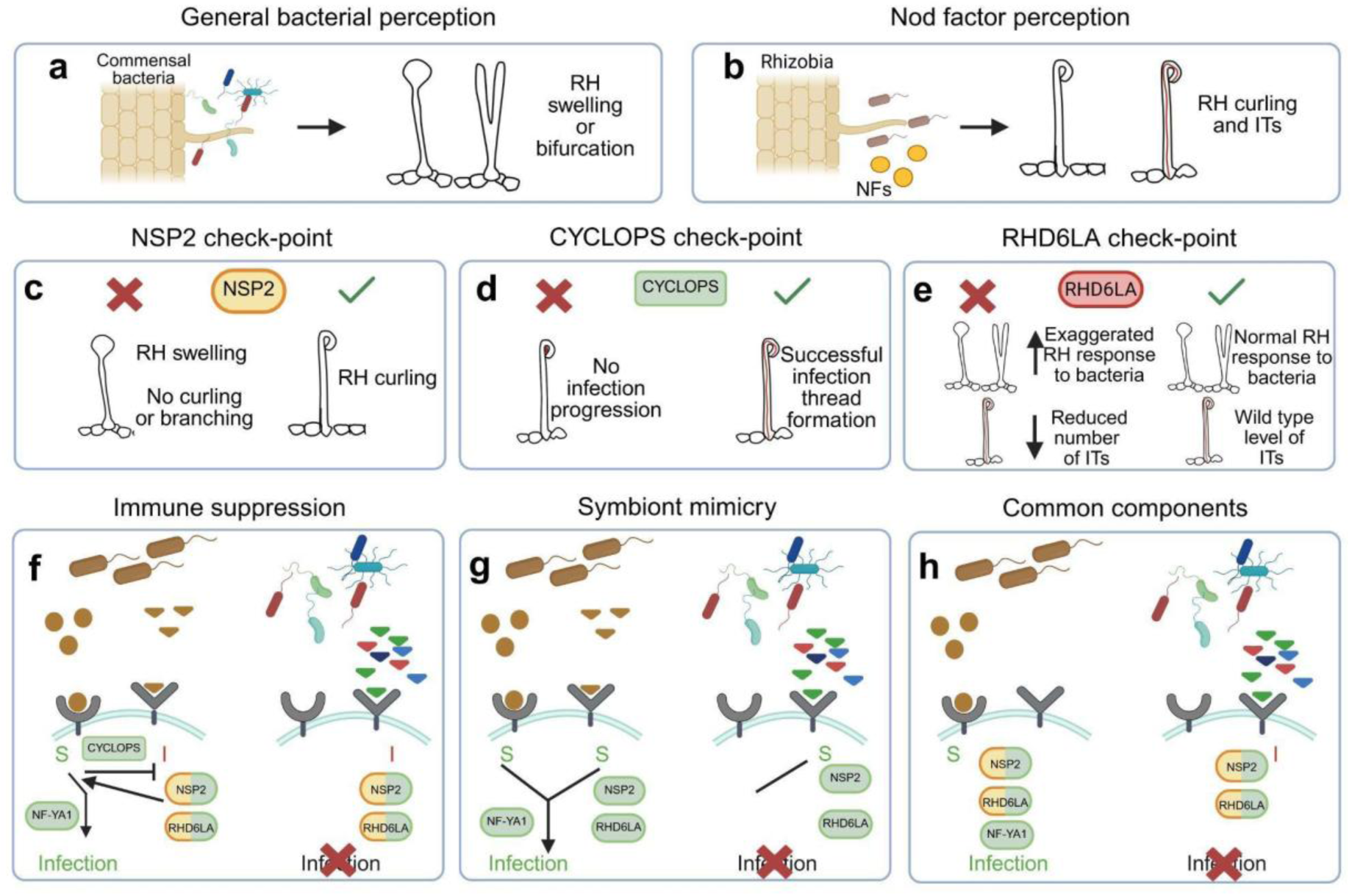
Scenarios for root hair responses to commensal and symbiotic bacteria. **a)** Root hair perception of commensal bacteria, which do not produce Nod factors, lead to swelling or bifurcation, but not to infection (Fig. 4 b-c). **b)** Nod factor-producing rhizobia induce root hair curling and infection thread formation. **c)** *nsp2* mutants respond to symbiotic rhizobia only with root hair swelling. No curling or infection thread formation is observed ^54^. **d)** Root hair curling is observed in *cyclops* mutants, but no infection threads are formed ^10^. **e)** *rhd6la* mutants show exaggerated root hair deformation in response to commensal bacteria and a reduced number of infection threads on rhizobium inoculation (Fig. 4 d-f). **f)** Immune suppression scenario. Symbiotic rhizobia induce both symbiotic (S) and immune (I) signalling. Symbiotic signalling suppresses immune signaling in a CYCLOPS-dependent manner. RHD6LA and NSP2 promote symbiont infection in combination with Nod factor signalling. Commensal bacteria only induce immune signalling. **g)** Symbiont mimicry scenario. The non-Nod factor signal is part of the symbiotic response. Commensal bacteria mimic the non-Nod factor signal, but lack of Nod factors prevents infection. **h)** Common components scenario. Symbiotic rhizobia do not necessarily induce immune signaling, but NSP2 and RHD6LA are downstream of both Nod factor and Nod factor-independent bacterial perception. Figure created with the assistance of BioRender.com.

This transcriptional overlap could in principle be caused by a general stress response in root hair cells that experience incompatible microbial interactions, or competition for nutrients, regardless of whether they result from challenge by non-symbiotic commensals or deficient CSSP signaling during rhizobium infection. If the overlap represented a general stress response, it would be expected to occur broadly across root hair cells exposed to commensals. However, the specificity of the transcriptional responses to rhizobium-responsive root hair argues against this hypothesis. Instead, it is likely that a general, Nod factor-independent, bacterial perception system exists in rhizobium-responsive root hairs alongside the specific Nod factor perception mediated by Nod factor receptors (**Fig. 5 f-h**).

Activation of this general bacterial perception system alone would not promote infection and could represent an immune signal, but the general system would also induce components, including RHD6LA and NSP2, that promote rhizobium infection in combination with Nod factor signalling (**Fig. 5d**). The fact that infection fails in *cyclops* rhizobium-responsive root hairs, despite initiation of Nod factor signalling, suggests that CYCLOPS could play a role in suppressing immune signaling (**Fig. 5f**).

Alternatively, the Nod factor-independent bacterial perception system could be an integral part of rhizobium recognition that has been hijacked by certain commensals, enabling them to activate part of the symbiotic program in responsive root hairs (**Fig. 5g**). By tagging along with Nod factor-producing rhizobia, this could allow commensals to gain access via intracellular infection to root nodules that make up attractive, nutrient-rich ecological niches. In this scenario, CYCLOPS would not be required for suppression of immunity, but merely for full activation of the symbiotic program to complement the Nod factor-independent signal.

A third option is that the transcriptional overlap is caused by common signaling components used downstream of both general bacterial and Nod factor perception (**Fig. 5h**). The transcriptional overlap between *cyclops* + R7A and wild-type + SynCom19 would then be due to Nod factor and general bacterial perception signals feeding into the same signalling pathway, which is normally pushed towards infection promotion by compatible rhizobia through CYCLOPS-dependent CSSP signalling.

Root hair morphological responses offer some support for this scenario. *nsp2* mutants are able to perceive Nod factors and initiate calcium spiking, but do not show root hair curling. Instead *nsp2* root hairs exhibit swelling and deformation reminiscent of the wild-type root hair response to commensals detailed here (**Fig 4. b-c, Fig. 5a,c**). This morphological root hair response to commensals is exaggerated in *rhd6la* mutants, which also show reduced infection thread and nodule counts (**Fig. 4-ef, Fig. 5e**). In contrast, Nod factor deficient rhizobia do not induce root hair deformation ^44^. So, whereas Nod factors are needed for root hair deformation in response to rhizobia, commensals can induce root hair deformation in the absence of Nod factor signalling, just like they can induce *NSP2* and *RHD6LA* expression independent of Nod factors.

The scenarios presented in **Fig. 5** are simplifications. In reality, Nod factors, conserved microbe-associated molecular patterns, microbial effectors and plant detection systems result in a great variation in interaction outcomes, especially when no fully compatible symbionts are available ^47^. This extends also to commensals co-colonising nodules with compatible rhizobia^48^. However, determining the reasons underlying the overlapping root hair transcriptional responses to commensals and symbionts could help us take the next steps in understanding how legumes balance the benefits of symbiosis with the risk of exploitation by potential pathogens.

From an evolutionary perspective, the most parsimonious explanation for our findings is that the general bacterial perception system in root hairs predates symbiosis-specific Nod factor perception, which would be a later adaptation added in the process of enabling bacterial endosymbiosis. However, the localisation of the transcriptional response to a subset of root hairs involved in rhizobium signalling would suggest that the general perception system in responsive root hairs could be a legume-specific innovation serving to stabilise rhizobium endosymbiosis by providing protection against cheater bacteria. The phylogenomic distribution of *RHD6LA* is compatible with this hypothesis, since *RHD6LA* appears to result from a legume-specific duplication occurring around the time intracellular rhizobium infection evolved and show signs of pseudogenization in peanut, which relies on intracellular infection (**Supplementary fig. 3**).

In conclusion, our study reveals a previously unappreciated complexity in plant-microbe interactions at the root hair interface (**Fig. 5**). The interplay between general bacterial perception and symbiosis-specific Nod factor signaling in select root hairs provides a new framework for investigating the molecular mechanisms underlying plant-microbe recognition and the evolution of symbiotic relationships.

## Methods

### Plant material

*Lotus japonicus* seeds from the Gifu accession (both wild type and mutants) were subjected to scarification using sandpaper, followed by sterilization using a 1% (v/v) sodium hypochlorite solution for 8-10 minutes. Subsequently, the seeds underwent five washes with sterile water in sterile conditions. After being kept at 4°C overnight, the seeds were moved to square Petri dishes for germination under a 16-hour day cycle (at 21 °C) followed by an 8-hour night cycle (at 19 °C). After four days the seeds with emerging radicles were moved to square plates containing 1.4% Agar Noble slopes supplemented with 0.25x B&D medium. These plates were covered with filter paper. A metal bar featuring 3-mm holes for roots was introduced at the upper edge of the agar slope. Plant growth plates containing 10 seedlings each were then inoculated with 500 µL of microbial communities suspended in water with an OD600 = 0.02 and applied along the length of the roots. For genetic studies, the *LORE1* lines 30107507 (*rhd6la-1*) and 30035826 (*rhd6la-2*) were used^41^ .

### Bacterial strains

The 19 bacterial strains (SynCom) were inoculated in tryptic soy broth (TSB) media for 2 days at 28°C, then centrifuged at 4,000 *g* for 15 min. The supernatant was discarded and the pellet was resuspended in 1 mL water. The OD600 was measured and the bacterial suspensions were diluted to obtain an OD600 equal to 0.02. The *M. loti* R7A rhizobia strain was cultured for 2 days at 28°C in yeast mannitol broth (YMB) agar plate, resuspended in water and diluted accordingly to obtain an OD600 equal to 0.02.

### Protoplast Isolation and scRNA-seq

For the isolation of protoplasts, whole roots were subjected to protoplasting under gentle agitation over a 3-hour period at room temperature using a 5 mL digestion solution. This digestion solution contained 10 mM MES (pH 5.7), 1.5% (w/v) cellulase R-10, 2% (w/v) macerozyme R-10, 0.4M D-sorbitol, 10 mM CaCl2, 5% (v/v) viscozyme, and 1% (w/v) BSA. The resultant intact protoplasts were separated by straining the protoplast-containing digestion solution through a 40 µM strainer into 15 mL Falcon tubes. This mixture was then combined with 5 mL of a 50% Optiprep solution, composed of 50% (v/v) Optiprep, 10 mM MES (pH 5.7), 0.4M D-sorbitol, 5mM KCl, and 10 mM CaCl2. To the combined solution, 2 mL of a 12.5% Optiprep solution (12.5% (v/v) Optiprep, 10 mM MES (pH 5.7), 0.4M D-sorbitol, 5mM KCl, 10 mM CaCl2) and 250 µL of a 0% Optiprep solution (10 mM MES (pH 5.7), 0.4M D-sorbitol, 5mM KCl, 10 mM CaCl2) were cautiously added in sequence. Subsequent to this, the tubes were centrifuged at 250*g* for 10 min at 4°C. Living protoplasts were collected at the interface between the 12.5% and 0% Optiprep solutions and counted using a Neubauer chamber. For the generation of single-cell RNA sequencing libraries, the Chromium Next GEM Single Cell 3’ Kit v3.1 (10X Genomics) was utilized in accordance with the manufacturer’s instructions, with the aim of recovering 5000 cells for each biological replicate.

### Raw data pre-processing, integration and clustering

The initial sequencing data were processed using Cell Ranger v6.1.2 from 10X Genomics. The genome assembly and gene annotations were established using *Lotus japonicus* Gifu v1.2 and Gifu v1.3 as references, respectively. These references are accessible via Lotus Base^31^ The aligner used for the analysis was STAR v2.7.2a ^49^. Employing the default parameters, "cellranger count" was executed. The subsequent stages utilized the "filtered_feature_bc_matrix" obtained from this process as input. Subsequent analyses were conducted using Seurat version 4.0.5 ^50^. Specifically, cells with an expression of fewer than 200 or more than 7500 genes, along with fewer than 500 UMIs, were removed from the dataset. Further filtering was performed based on the expression of genes associated with mitochondrial and chloroplast genomes. Cells expressing less than 5% of their total read counts were retained after this filtering step. Normalization across all samples was achieved using the "sctransform" function embedded within Seurat, where the "vars.to.regress" parameter was set to encompass mitochondrial and chloroplast genes. Subsequently, the samples were integrated through Seurat’s canonical correlation analysis integration pipeline. Employing the integrated data assay, a Principal Component Analysis (PCA) was performed using Seurat’s default function for dimensionality reduction. Following this, the "FindNeighbors" and "RunUMAP" functions were applied, utilizing 50 principal components for the entire datasets and 30 for the subsets related to root hairs. Cells were clustered using an unsupervised Louvain clustering algorithm with the default resolution of 0.8.

For the integrated object containing data from the current study and 5 and 10 dpi *M. loti* R7A data^32^, analysis was performed using Seurat 5.1.0. The same analysis procedure as described above was followed, except for the Louvain clustering resolution, where a value of 0.5 was used.

The topometry analysis ^34^ was performed on the scaled expressions for the 3000 genes in the integrated assay of the Seurat object. The analysis was executed using the default command with the kernel set to ’bw_adaptive’, the eigen methods used were ’msDM’, ’DM’ and the ’MAP’ projection. The clustering resolution was set to 2.

ShinyCell was used to create the web interface for browsing the single-cell datasets (https://lotussinglecell.shinyapps.io/lotus_japonicus_single_cell_microbiome/) ^51^

### Differential gene expression

Differential gene expression analysis between treatments for each cluster and the markers for specific groups of cells were identified using the "MAST" algorithm v 1.16 61 with the “min.pct” = 0.01 parameter. Genes satisfying the criteria of an adjusted p-value of ≤ 0.05 and a log fold change greater than 0.25 were classified as differentially expressed.

### Correlation analysis

We identified differentially expressed genes in the three cell populations (Infected_RH_5dpi, SynCom19_RHsubcluster7, and RH_*cyclops*) compared to the root hairs for their corresponding mock sample using the default method in Seurat. Genes were filtered for pct.2 ≤ 0.1 and adjusted p-value ≤ 0.05. We computed log2 average pseudobulk counts for all the differentially expressed genes across the three cell populations. The correlogram was constructed using the psych R package ^52^. The significance between correlation values was calculated using the cocor package with the Pearson and Filon’s (1898) z test ^53^.

### Confocal microscopy

Confocal microscopy was performed with Zeiss LSM780 microscope.

### RHD6LA Phylogenetic tree

Protein sequences were retrieved from Phytozome 13 (The plant genomics resource). Protein sequences have been aligned with MUSCLE software and a Maximum Likelihood phylogenetic tree has been generated using 500 bootstrap replicates for node support values (WAG model, Gamma distributed, Partial deletion 90).

### Data availability

The sequencing data generated in this study have been deposited as ENA project accession PRJEB89066. UMAPs and gene expression data can be browsed at [https://lotussinglecell.shinyapps.io/lotus_japonicus_single_cell_microbiome/]. All generated gene lists are provided as Source Data. Source data are provided with this paper.

## Author contributions

F.T., J.Q. and S.U.A. conceived and designed experiments; F.T. and J.Q. conducted experiments; F.T. and L.I.F. performed the bioinformatic scRNAseq analysis, L.I.F. developed the Shiny app, F.T., J.Q. and S.U.A. analyzed the data; S.J.V.C genotyped the mutant alleles; J.Q. and F.T. performed microscopy; F.T. drafted the first version of the manuscript; J.Q. and S.U.A. edited the manuscript with input from all authors.

## Acknowledgements

This work was supported by the Novo Nordisk Foundation grant [no. 307NNF19SA0059362] for the InRoot project coordinated by Jens Stougaard and by Independent Research Fund Denmark (grant agreement no. 1026-00032B) to S.U.A.

