## Supplemental Figure 1, Supplemental Figure 2, Supplemental Figure 3 for "RHD6LA regulates root hair responses to both symbionts and commensals"

### Supplementary Figures

a

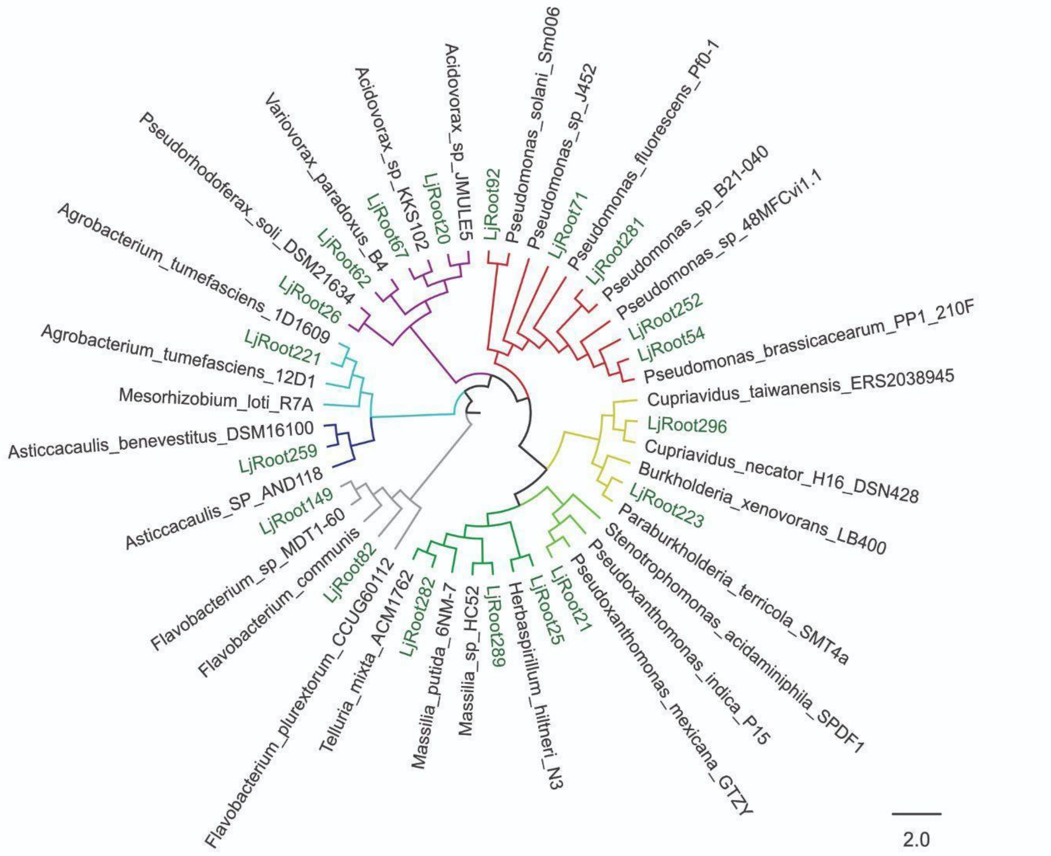


b

| Isolate name | Family | Genus | Species (>95% ANI) | Closest reference (>90% ANI) |
|---|---|---|---|---|
| LjRoot223 | Burkholderiaceae | Paraburkholderia | *P. terricola* | *P. terricola SMT4a* |
| LjRoot296 | Burkholderiaceae | Cupriavidus | *C. necator* | *C. necator H16* |
| LjRoot259 | Caulobacteraceae | Asticcacaulis | - | - |
| LjRoot20 | Comamonadaceae | Acidovorax | - | *A. sp. JMULE5* |
| LjRoot26 | Comamonadaceae | Pseudorhodoferax | - | *P. soli DSM21634* |
| LjRoot62 | Comamonadaceae | Variovorax | - | *V. paradoxus B4* |
| LjRoot67 | Comamonadaceae | Acidovorax | - | - |
| LjRoot149 | Flavobacteriaceae | Flavobacterium | - | - |
| LjRoot82 | Flavobacteriaceae | Flavobacterium | - | *F. plurextorum CCUG 60112* |
| LjRoot25 | Oxalobacteraceae | Herbaspirillum | - | *H. hiltneri N3* |
| LjRoot282 | Oxalobacteraceae | Telluria | - | *Telluria mixta ACM1762* |
| LjRoot289 | Oxalobacteraceae | Massilia | - | - |
| LjRoot252 | Pseudomonadaceae | Pseudomonas | *P. brassicacearum* | *P. brassicacearum PP1_210F* |
| LjRoot281 | Pseudomonadaceae | Pseudomonas | *P. sp. B21-040* | *P. sp. B21-040* |
| LjRoot54 | Pseudomonadaceae | Pseudomonas | *P. brassicacearum* | *P. brassicacearum PP1_210F* |
| LjRoot71 | Pseudomonadaceae | Pseudomonas | - | - |
| LjRoot92 | Pseudomonadaceae | Pseudomonas | *P. solani* | *P. solani Sm006* |
| LjRoot221 | Rhizobiaceae | Agrobacterium | *A. tumefaciens* | *A. tumefaciens 1D1609* |
| LjRoot21 | Xanthomonanadaceae | Pseudoxanthomonas | - | - |
| R7A | Rhizobiaceae | Mesorhizobium | *M. loti* | *M. loti R7A* |

**Supplementary figure 1**. **Bacterial strains used to create the SynCom19 inoculum for the scRNA-Seq experiment.** a) phylogenetic tree generated with recA full length DNA sequences. b) List of the isolates and taxonomies

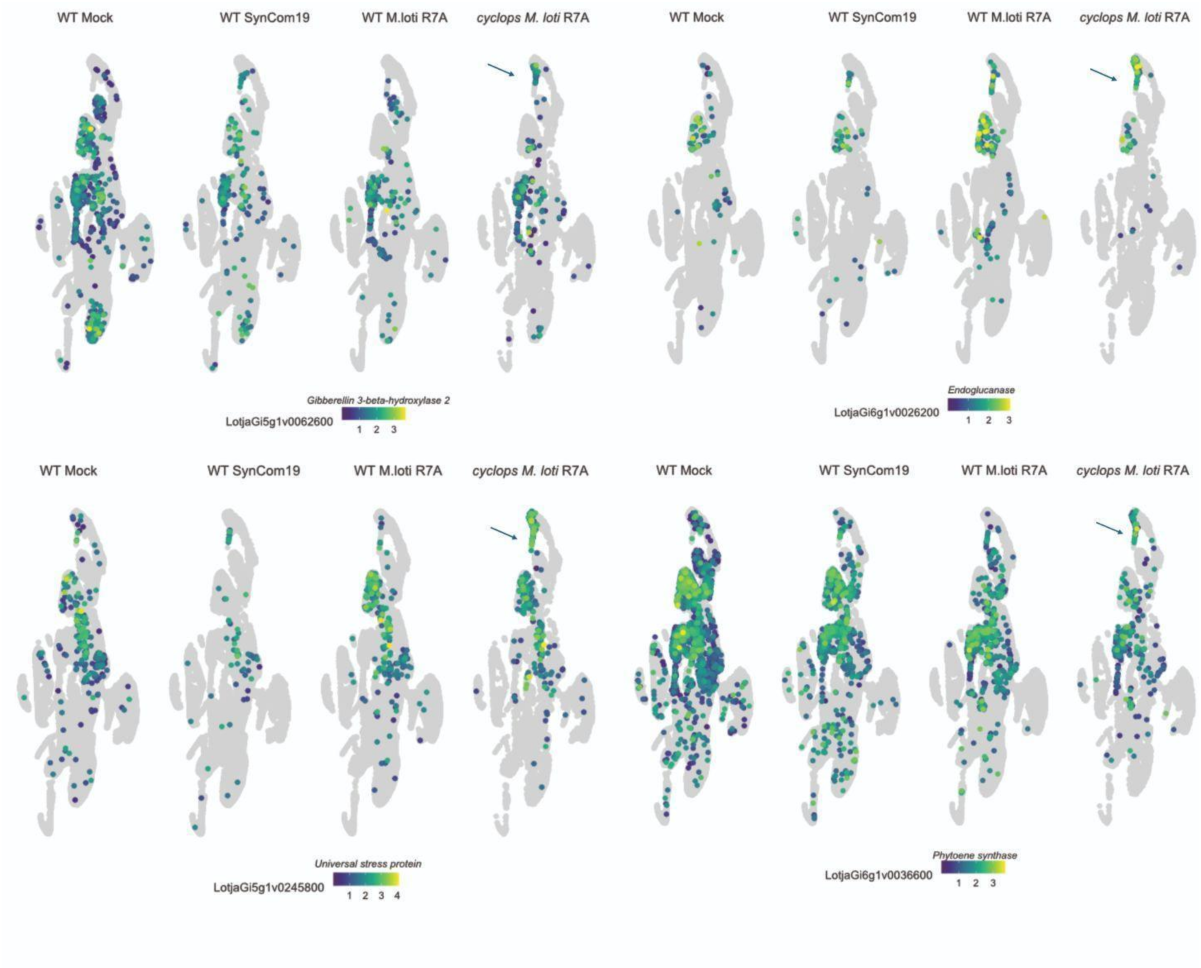


**Supplementary figure 2.** Expression patterns of selected genes commonly induced in a specific root hair population in WT SynCom19 and *cyclops* M.loti R7A

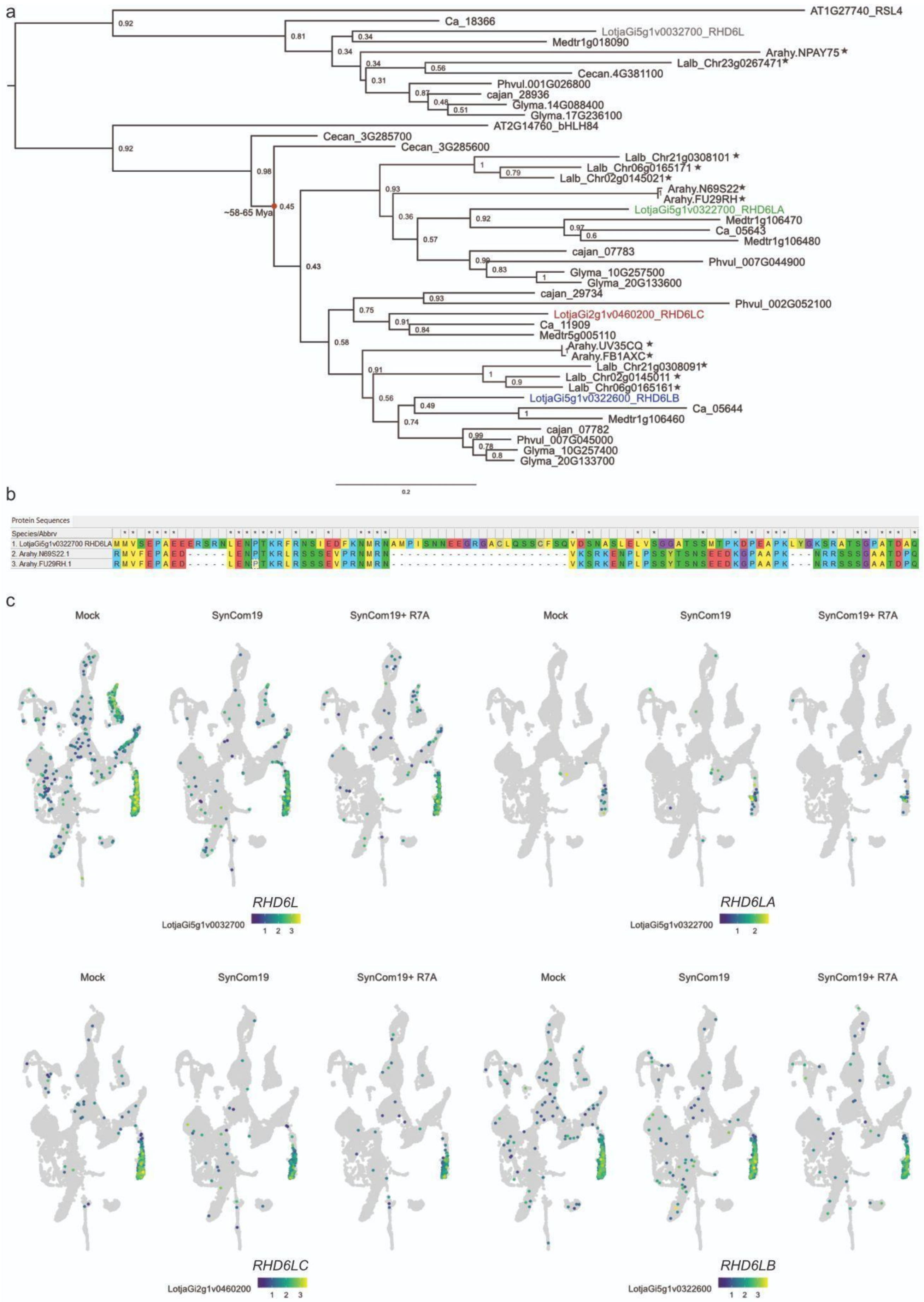
a
AT1G27740_RSL4
Ca_18366
LotjaGi5g1v0032700_RHD6L
Medtr1g018090
Arahy.NPAY75★
Lalb_Chr23g0267471★
Cecan.4G381100
Phvul.001G026800
cajan_28936
Glyma.14G088400
Glyma.17G236100
AT2G14760_bHLH84
Cecan_3G285700
Cecan_3G285600
~58-65 Mya
Lalb_Chr21g0308101★
Lalb_Chr06g0165171★
Lalb_Chr02g0145021★
Arahy.N69S22★
Arahy.FU29RH★
LotjaGi5g1v0322700_RHD6LA
Medtr1g106470
Ca_05643
Medtr1g106480
cajan_07783
Phvul_007G044900
Glyma_10G257500
Glyma_20G133600
cajan_29734
Phvul_002G052100
LotjaGi2g1v0460200_RHD6LC
Ca_11909
Medtr5g005110
Arahy.UV35CQ★
Arahy.FB1AXC★
Lalb_Chr21g0308091★
Lalb_Chr02g0145011★
Lalb_Chr06g0165161★
LotjaGi5g1v0322600_RHD6LB
Ca_05644
Medtr1g106460
cajan_07782
Phvul_007G045000
Glyma_10G257400
Glyma_20G133700
0.2
b
Protein Sequences
Species/Abbrv
1. LotjaGi5g1v0322700 RHD6LA
2. Arahy.N69S22.1
3. Arahy.FU29RH.1
c
Mock
SynCom19
SynCom19+ R7A
RHD6L
LotjaGi5g1v0032700
RHD6LA
LotjaGi5g1v0322700
RHD6LC
LotjaGi2g1v0460200
RHD6LB
LotjaGi5g1v0322600

**Supplementary Figure 3. Evolution and expression of the RHD6L family. a)** Maximum Likelihood phylogenetic tree (500 bootstrap replicates; node support values shown) based on protein sequence alignments generated with MUSCLE. *Lotus japonicus* proteins are highlighted in distinct colors. Stars indicate species known to perform intercellular or hybrid infection processes (*Arachis hypogaea*, *Lupinus albus*). *Cercis canadensis* (Ceca) is shown as an early-diverging lineage marking the split between Cercidoideae and Faboideae, estimated at 58–65 million years ago (red circle). **b)** Protein sequence alignment snapshot (LjRHD6LA, Arahy.RHD6LA_copy1, Arahy.RHD6LA_copy2) showing three conserved gaps located within the core of the "Transcription Factor bHLH83-related" domain (PTHR16223; amino acids 81–309). **c)** UMAP plots illustrating single-cell expression patterns of the four *L. japonicus* RHD6L-like paralogs. The ancestral *RHD6L* is broadly expressed across tissues, with enrichment in root hairs (RH) and root caps. *RHD6LA* shows specific expression in a subset of RH cells upon SynCom19 inoculation. *RHD6LB* and *RHD6LC* display strong expressions restricted to RHs.
